# Somatostatin-positive Interneurons Contribute to Seizures in *SCN8A* Epileptic Encephalopathy

**DOI:** 10.1101/2021.02.05.429987

**Authors:** Eric R. Wengert, Kyle C.A. Wedgwood, Pravin K. Wagley, Samantha M. Strohm, Payal S. Panchal, Abrar Idrissi Majidi, Ian C. Wenker, Ronald P. Gaykema, Manoj K. Patel

## Abstract

*SCN8A* epileptic encephalopathy is a devastating epilepsy syndrome caused by mutant *SCN8A* which encodes the voltage-gated sodium channel Na_V_1.6. To date, it is unclear if and how inhibitory interneurons, which express Na_V_1.6, influence disease pathology. We found that selective expression of the R1872W mutation in somatostatin (SST) interneurons was sufficient to convey susceptibility to audiogenic seizures. SST interneurons from mutant mice were hyperexcitable but hypersensitive to action potential failure via depolarization block under normal and seizure-like conditions. Remarkably, GqDREADD-mediated activation of wild-type SST interneurons resulted in prolonged electrographic seizures and was accompanied by SST hyperexcitability and depolarization block. Aberrantly large persistent sodium currents, a hallmark of *SCN8A* mutations, were observed and were found to contribute directly to aberrant SST physiology in computational and pharmacological experiments. These novel findings demonstrate a critical and previously unidentified contribution of SST interneurons to seizure generation not only in *SCN8A* encephalopathy, but epilepsy in general.

## Introduction

*SCN8A* encephalopathy is a severe genetic epilepsy syndrome caused by *de novo* mutations in the *SCN8A* gene which codes for the voltage-gated sodium channel isoform Na_V_1.6 (Gardella et al., 2018; Larsen et al., 2015; Ohba et al., 2014; Veeramah et al., 2012). Patients exhibit a panoply of devastating symptoms including refractory epilepsy, cognitive impairment, motor dysfunction, sensorineural hearing loss, and have a substantial risk of sudden unexpected death in epilepsy (SUDEP) (Gardella et al., 2018; Larsen et al., 2015; Ohba et al., 2014; Veeramah et al., 2012).

Previous studies using patient-derived *SCN8A* mutations have sought to mechanistically understand how these *de novo* mutations alter biophysical properties of voltage-gated sodium channels, intrinsic excitability of single neurons, and network dynamics underlying behavioral seizures (Baker et al., 2018; Bunton-Stasyshyn et al., 2019; Estacion et al., 2014; de Kovel et al., 2014; Lopez-Santiago et al., 2017; Ottolini et al., 2017; Patel et al., 2016; Wagnon et al., 2016; Wengert et al., 2019a). Understanding the circuit-basis for *SCN8A* encephalopathy requires evaluation of the various cellular subpopulations which comprise neural circuits. Although Na_V_1.6 is expressed in inhibitory interneurons (Li et al., 2014; Lorincz and Nusser, 2008; Makinson et al., 2017), the mechanisms of network dysfunction in *SCN8A* encephalopathy is greatly limited by the current lack of understanding how inhibitory interneuron function is affected by the expression of mutant Na_V_1.6. Inhibitory interneuron function is critical for constraining overall network excitability and preventing seizures (Bernard et al., 2000; Kumar and Buckmaster, 2006; Markram et al., 2004). Dysfunction of inhibitory interneurons has therefore been proposed as an important mechanism of epilepsy (Bernard et al., 2000; Stafstrom, 2013). With respect to sodium channel epileptic encephalopathies, hypofunction of inhibitory interneurons has typically been associated with loss-of-function *SCN1A* mutations which result in Dravet Syndrome (Cheah et al., 2012; Escayg and Goldin, 2010; Rhodes et al., 2004; Tai et al., 2014). Previous studies utilizing mouse models of Dravet Syndrome have confirmed that expression of the mutant *SCN1A* allele in GABAergic interneurons is sufficient to recapitulate the key phenotypes of Dravet Syndrome (Cheah et al., 2012). Measurements of intrinsic excitability also revealed impaired excitability of parvalbumin-positive (PV) and somatostatin-positive (SST) inhibitory interneurons at certain key points in development (Favero et al., 2018; Tai et al., 2014). The phenotypic severity of Dravet Syndrome suggests that seemingly small changes to inhibitory interneuron excitability (reduction of max firing frequency by ∼20%) may have profound effects for overall network excitability and seizure behavior (Tai et al., 2014).

In this study, we utilized two mouse models of patient derived *SCN8A* mutations; N1768D (*Scn8a*^*D*/+^) which allows global heterozygous expression and a conditional mouse in which the R1872W (*Scn8a*^*w*/+^) mutation is dependent on Cre recombinase. Mice globally expressing either mutation are susceptible to audiogenic seizures, providing a powerful experimental approach to assess seizure susceptibility (manuscript in review). Using these two clinically relevant mouse models of epileptic encephalopathy, we report that specific expression of R1872W in SST interneurons (*Scn8a*-SST^W/+^), but not expression in forebrain excitatory neurons (*Scn8a*-EMX1^W/+^), results in audiogenic seizures, indicating that abnormal activity of SST interneurons uniquely contributes to seizures in *SCN8A* encephalopathy. In *Scn8a*^*D*/+^ mice and mice specifically expressing R1872W in SST interneurons (*Scn8a*-SST^W/+^), recordings from SST interneurons showed an initial steady-state hyperexcitability, followed by action potential failure via depolarization block. Dual-cell recordings revealed that mutant SST interneurons are particularly sensitive to depolarization block under seizure-like conditions and that depolarization block occurs coincidently with pyramidal neuron ictal discharges. Counterintuitively, GqDREADD-mediated chemogenetic activation of wild-type control SST interneurons was sufficient to facilitate initial hyperexcitability followed by depolarization block and *status epilepticus* in the mice, directly demonstrating the proconvulsant effect of aberrant SST physiology. Recordings of voltage-gated sodium channel activity identified elevated persistent sodium currents (I_NaP_) in mutant SST interneurons. Computational modelling revealed that elevating I_NaP_ was singly sufficient to induce initial hyperexcitability and premature entry into depolarization block. Lastly, we validated the model predictions by demonstrating that application of veratridine, a pharmacological activator of I_NaP_, to wild-type SST interneurons induces premature depolarization block. Overall, our results reveal a previously unappreciated mechanism by which SST interneurons contribute to seizures in *SCN8A* encephalopathy and may potentially generate novel therapeutic intervention strategies through targeting this important neuronal subpopulation.

## Results

Using a model of *SCN8A* encephalopathy in which the patient-derived mutation R1872W is expressed in a Cre-dependent manner, we sought to determine the cell-type specific contribution to audiogenic seizure susceptibility. Acoustic stimulation of mice globally-expressing the R1872W *Scn8a* mutation (*Scn8a*-EIIa^W/+^) caused audiogenic seizures (7 of 8 mice; 87.5%; Figure 1A-C). Seizures were characterized by a period of wild-running, a tonic phase associated with hindlimb extension, and sudden death (Figure 1A; Video 1). To our surprise, expression of the R1872W *Scn8a* mutation in forebrain excitatory neurons (*Scn8a*-EMX1^W/+^) did not reliably convey susceptibility to audiogenic seizures, with only 1 of 9 mice exhibiting a single brief bout of wild-running; (Video 2; ^***^P<0.01 Fisher’s Exact test; Figure 1B, C, E). In contrast, expression of the R1872W *Scn8a* mutation selectively in somatostatin-positive (SST) inhibitory interneurons (*Scn8a*-SST^W/+^) was sufficient to induce an audiogenic seizure in 18 of 22 (∼82%), typified by a wild-running phase, a clonic phase characterized by convulsions and jerking of limbs, followed by recovery (Video 3; Figure 1A-C). Audiogenic seizures were never observed in WT littermate control mice (n=26: Video 4). These results indicate that expression of R1872W specifically in SST interneurons is sufficient to induce audiogenic seizures.

**Figure 1.**
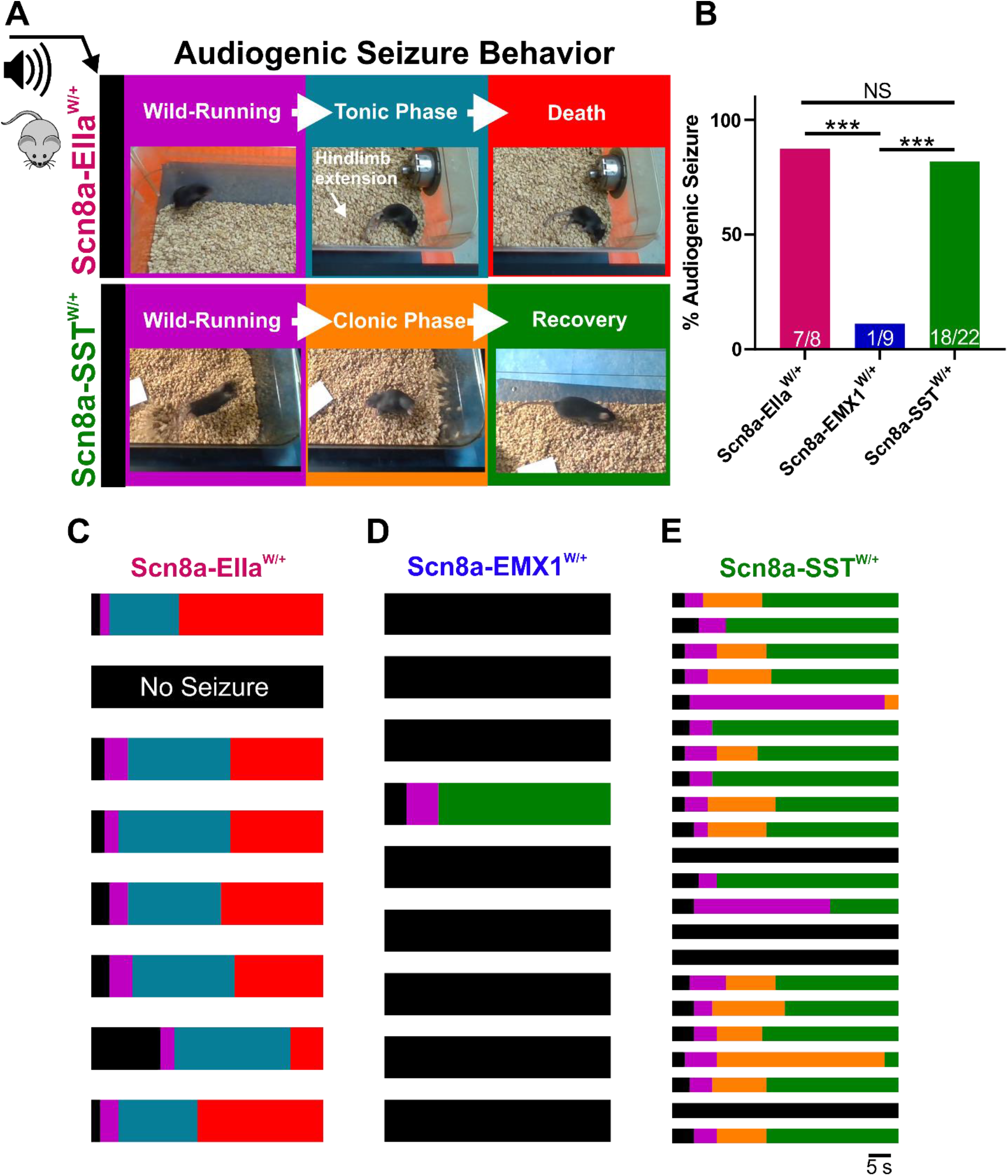
Somatostatin interneuron-specific expression of mutant *Scn8a* is sufficient for susceptibility to audiogenic seizures. **A**. Audiogenic seizure behavior in mice with cell-type specific expression of R1872W *Scn8a* mutation. Upon high-intensity acoustic stimulation, *Scn8a*-EIIa^W/+^ mice exhibit wild-running (purple) followed by a tonic phase characterized by hindlimb extension (blue) which is followed by collapse of breathing and death (red). *Scn8a*-SST^W/+^ mice exhibit wild-running (purple), but progress to a convulsive clonic phase characterized by repetitive shaking and limb-jerking (orange) followed by recovery (green). **B**. Propensity of audiogenic seizures in *Scn8a*-EIIa^W/+^ (magenta: 87.5 %), *Scn8a*-EMX1^W/+^ (∼11%) and *Scn8a*-SST^W/+^ (green: ∼82%) mice. Significance determined by Fisher’s exact test (^*^P<0.01, ^***^P<0.001). **C-E**. Color-coded raster plots for seizure behavior of individual *Scn8a*-EIIa^W/+^ (C), *Scn8a*-EMX1^W/+^ (D), and *Scn8a*-SST^W/+^ (E) mice.

We next sought to determine the impact of N1768D and R1872W patient-derived *SCN8A* mutations on SST interneuron function with a goal to better understand how alterations in SST interneuron excitability could lead to seizure susceptibility. We conducted whole-cell patch-clamp electrophysiology recordings of SST interneurons from layer V somatosensory cortex from adult *Scn8a*^*D*/+^, *Scn8a*-SST^W/+^ and littermate WT mice (Figure 2A). Relative to WT controls, a greater proportion of SST interneurons from *Scn8a*^*D*/+^ and *Scn8a*-SST^W/+^ were spontaneously active (^***^P<0.01; Fisher’s Exact test; Figure 2B-E) each with significantly greater frequency of spontaneous action potentials (APs) (^*^P<0.05 and ^**^P<0.01 relative to WT; Kruskal-Wallis test followed by Dunn’s test; Figure 2F). Analysis of membrane and AP properties revealed that, relative to WT, both *Scn8a*^*D*/+^ and *Scn8a*-SST^W/+^ SST interneurons had depolarized resting membrane potentials and decreased rheobases likely contributing to their steady-state hyperexcitability (Table 1). Additionally, *Scn8a*^*D*/+^ SST interneurons had an elevated input resistance and *Scn8a*-SST^W/+^ interneurons had hyperpolarized AP thresholds and increased AP duration (APD_50_; Table 1).

**Table 1:** Membrane and action potential properties of WT, *Scn8a*^*D*/+^, and *Scn8a*-SST^W/+^ somatostatin inhibitory interneurons.

|  | V<sub>m</sub> (mV) | AP Threshold (mV) | Upstroke Velocity (mV/ms) | Downstroke Velocity (mV/ms) | AP Amplitude (mV) | APD<sub>50</sub> (ms) | Input Resistance (MΩ) | Rheobase (pA) |
| --- | --- | --- | --- | --- | --- | --- | --- | --- |
| WT (n=32) | -60.7 ± 1.7 | -36.8 ± 0.8 | 323 ± 16 | -157 ± 13 | 74.0 ± 2.0 | 0.62 ± 0.03 | 215 ± 18 | 82 ± 12 |
| *Scn8a*<sup>D/+</sup> (n=33) | -52.9 ± 1.5** | -38.4 ± 0.8 | 290 ± 18 | -131 ± 7 | 76.7 ± 1.4 | 0.69 ± 0.03 | 298 ± 20* | 37 ± 7* |
| *Scn8a*-SST<sup>W/+</sup> (n=31) | -54.4 ± 1.6* | -40.84 ± 1.2** | 231 ± 2.8*** | -87 ± 5.7*** | 73.9 ± 2.0 | 1.0 ± 0.06*** | 250 ± 15 | 37 ± 5* |

Values are mean ± SEM. \*P<0.05; \*\*P<0.01, \*\*\*P<0.001 compared to WT using one-way ANOVA followed by Dunnett's multiple comparisons test.

**Figure 2.**
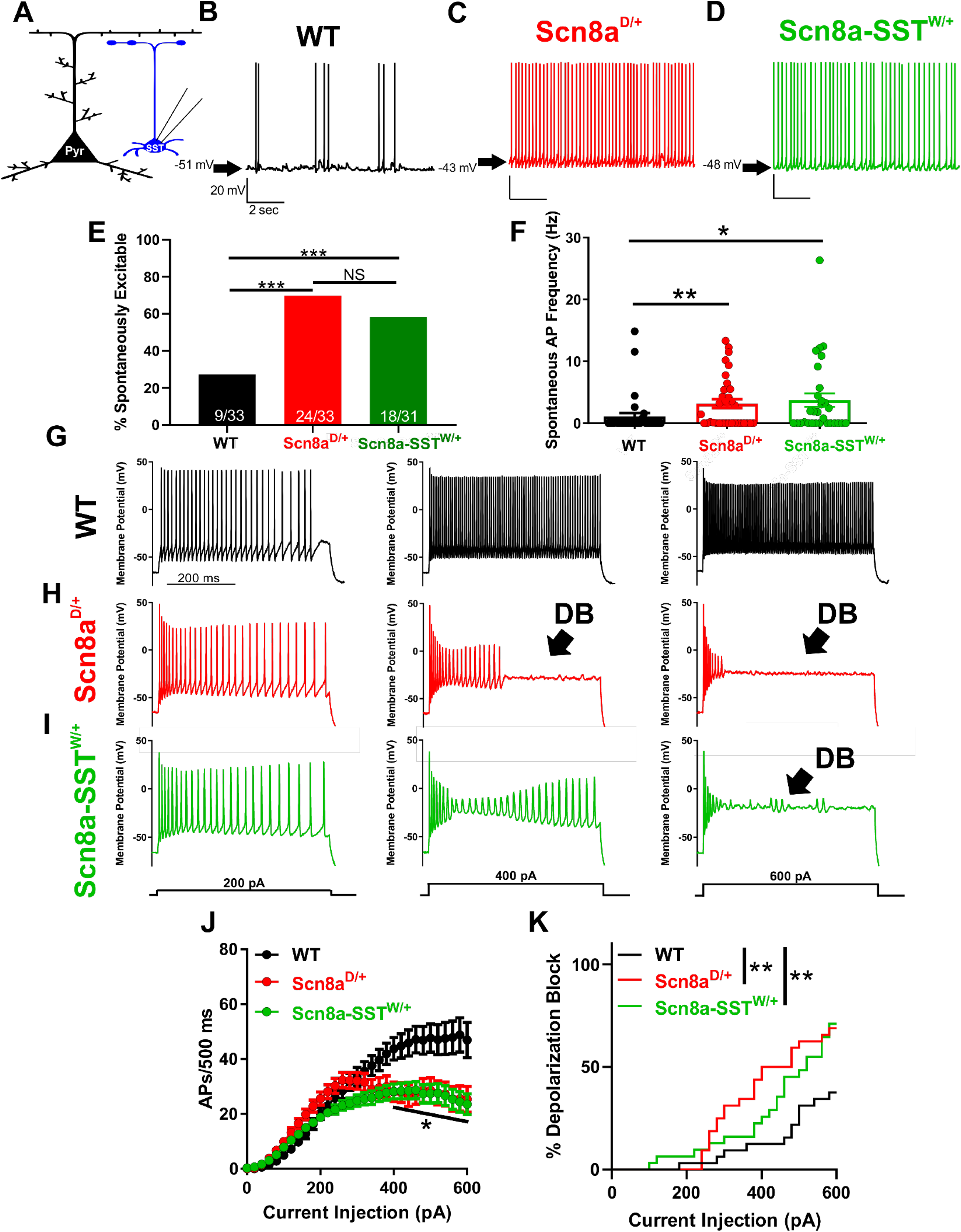
*Scn8a*^*D*/+^ and *Scn8a*-SST^W/+^ somatostatin interneurons are hyperexcitable and readily enter depolarization block. **A**. Whole-cell recordings collected from WT, *Scn8a*^*D*/+^, and *Scn8a*-SST^W/+^ somatosensory layer V SST interneurons (blue). **B-D**. Representative example traces of spontaneous excitability of WT (B; black), *Scn8a*^*D*/+^ (C; red) and *Scn8a*-SST^W/+^ (D; green) SST interneurons. Arrows indicate membrane potential between spontaneous APs. **E**. Only 9 of 33 (∼27%) WT SST interneurons were spontaneously excitable, whereas 24 of 33 (∼73%) *Scn8a*^*D*/+^ and 18 of 31 (∼58%) *Scn8a*-SST^W/+^ spontaneously fired APs (^***^P<0.01 by Fisher’s exact test). **F**. Average spontaneous firing frequencies for WT (black), *Scn8a*^*D*/+^ (red), and *Scn8a*-SST^W/+^ (green) SST interneurons. Statistical significance calculated by Kruskal-Wallis test followed by Dunn’s multiple comparisons (^*^P<0.05; ^**^P<0.01). **G-I**. Representative traces for WT (G; black), *Scn8a*^*D*/+^ (H; red), and *Scn8a*-SST^W/+^ (I; green) SST interneurons eliciting APs in response to 500-ms current injections of 200, 400, and 600 pA. Depolarization block is indicated (arrow; DB) in the *Scn8a*^*D*/+^ (H; red) and *Scn8a*-SST^W/+^ (I; green) interneurons. **J**. Average number of APs elicited relative to current injection magnitude for WT (black; n=33, 7 mice), *Scn8a*^*D*/+^ (red; n=33, 6 mice), and *Scn8a*-SST^W/+^ (green; n=31, 4 mice). At current injections >400 pA, both *Scn8a*^*D*/+^ and *Scn8a*-SST^W/+^ AP frequencies were reduced relative to WT (^*^P<0.05; Two-way ANOVA followed by Tukey’s correction for multiple comparisons). **K**. Cumulative distribution of SST interneuron entry into depolarization block relative to current injection magnitude for WT (black), *Scn8a*^*D*/+^ (red), and *Scn8a*-SST^W/+^ (green) mice. Curve comparison by Log-rank Mantel-Cox test (^**^P<0.01).

We subjected each neuron to a range of depolarizing current injections to characterize intrinsic excitability. Relative to WT controls (n=33, 7 mice), SST interneurons from both *Scn8a*^*D*/+^ (n=33, 6 mice) and *Scn8a*-SST^W/+^ mice (n=31, 4 mice) exhibited progressive AP failure at current injections above 400 pA as the interneurons entered depolarization block: At 600 pA, WT SST interneurons fired 47 ± 6 APs/500 ms (94 Hz), *Scn8a*^*D*/+^ fired 25 ± 5 APs/500 ms (50 Hz), and *Scn8a*-SST^W/+^ fired 23 ± 4 APs/500 ms (46 Hz; ^***^P<0.001 Two-way ANOVA with Tukey’s correction for multiple comparisons; Figure 2G-J). Over the range of current injection magnitudes, both *Scn8a*^*D*/+^ and *Scn8a*-SST^W/+^ interneurons were significantly more prone to depolarization block than WT counterparts (^**^P<0.01; Log-rank Mantel-Cox test; Figure 2K).

Prior studies have described cellular interplay between excitatory pyramidal neurons and fast-spiking interneurons during brain-slice seizure-like activity and *in vivo* seizures in which interneuron depolarization block occurs coincidentally with pyramidal neuron ictal discharges (Cammarota et al., 2013; Jayant et al., 2019; Parrish et al., 2019; Swadlow, 2003; Ziburkus et al., 2006). To test whether SST interneurons expressing *Scn8a* mutations also exhibit depolarization block seizure-like events coincident with pyramidal ictal discharges, we simultaneously recorded from a SST interneuron and a nearby pyramidal neuron and induced seizure-like activity by applying a bath solution with 0 Mg^2+^ and 4-AP (50 μM) (Ziburkus et al., 2006). Under these conditions, although neurons in all groups increased their excitability, we observed a significantly greater propensity for *Scn8a*^*D*/+^ (n= 12, 3 mice; Figure 3B) and *Scn8a*-SST^W/+^ (n=10, 3 mice; Figure 3C) SST interneurons to spontaneously exhibit depolarization block events compared with WT SST interneurons (n=9, 3 mice; Figure 3A) supporting our findings that expression of mutant *SCN8A* renders SST interneurons prone to depolarization block (^**^P<0.01 Log-rank Mantel-Cox test; Figure 3A-C, E). Consistent with previous reports in fast-spiking interneurons (Cammarota et al., 2013; Jayant et al., 2019; Parrish et al., 2019; Swadlow, 2003; Ziburkus et al., 2006), instances of SST interneuron depolarization block were entirely concomitant with seizure-like ictal discharges of nearby layer V pyramidal neurons (Figure 3B-D). Only when the bath concentration of the proconvulsant 4-AP was increased to 100 μM did we observe similar ictal-discharges of pyramidal neurons and depolarization block events in WT SST interneurons (Figure 3D-E). These results demonstrate not only that mutant SST interneurons are hypersensitive to depolarization block in the seizure-like context, but also that simultaneous SST depolarization block is coincident with pyramidal neuron bursting activity, a hallmark of seizure-like activity.

**Figure 3.**
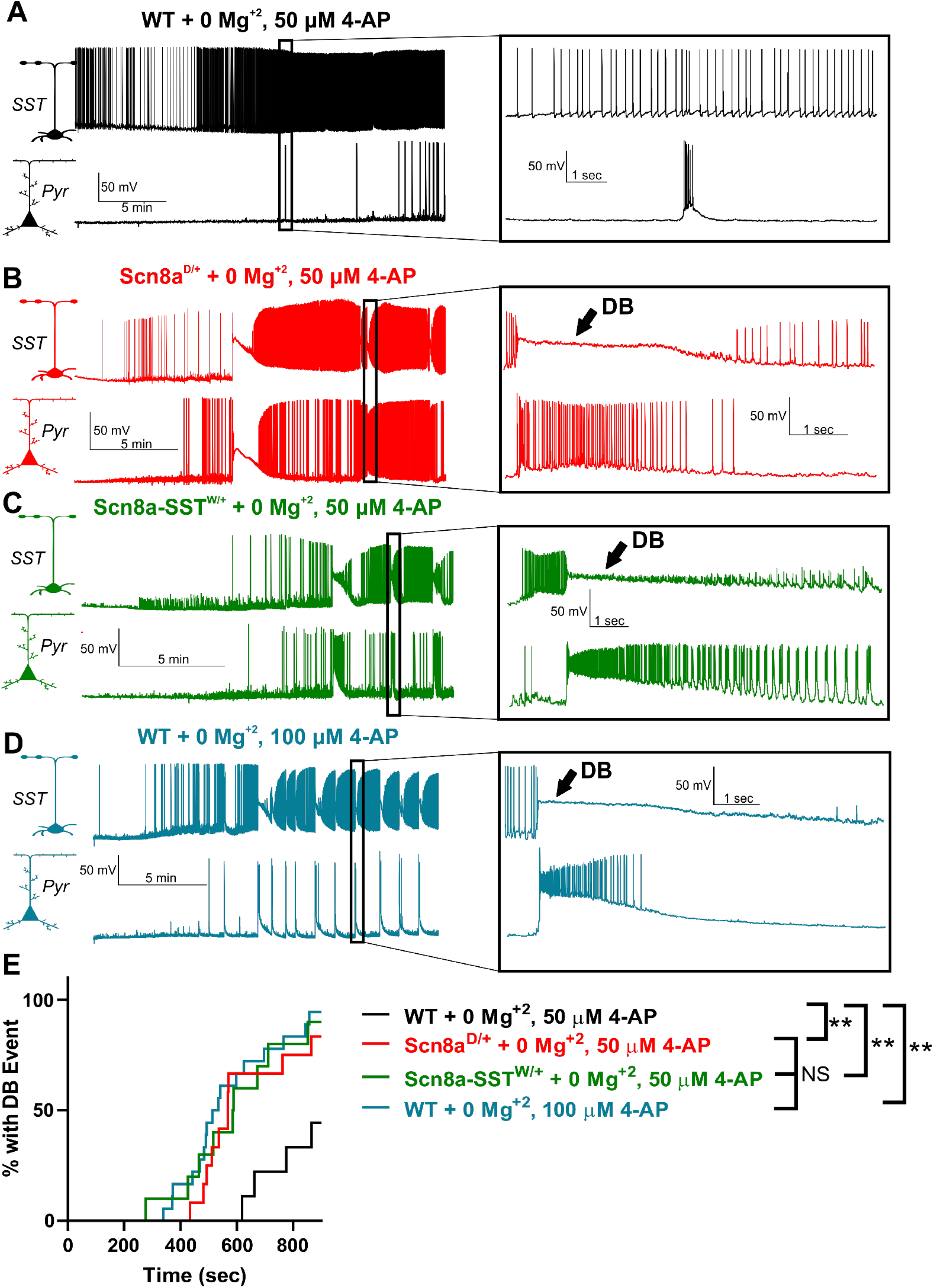
Hypersensitivity of mutant SST interneurons to depolarization block of somatostatin during seizure-like activity. **A-D**. Representative example traces of WT (A; black), *Scn8a*^*D*/+^ (B; red), and *Scn8a*-SST^W/+^ (C; green) SST-pyramidal neuron pairs exposed to bath solution containing 0 Mg^2+^, 50 μM 4-AP (A-C) and WT SST-pyramidal neuron pairs exposed to 0 Mg^2+^, 100 μM 4-AP (D). Expanded views indicate example synchronous SST interneuron depolarization block and pyramidal neuron ictal discharge events (B-D) which were not present in most neuronal pairs in the control group (A). **E**. Cumulative distribution plot of seizure-like events over time reveals that *Scn8a*^*D*/+^ + 0 Mg^+2^; 50 μM (red), *Scn8a*-SST^W/+^ + 0 Mg^+2^; 50 μM (green), and WT + 0 Mg^+2^; 100 μM (blue) were hypersensitive to depolarization block-mediated seizure-like events relative to WT + 0 Mg^+2^; 50 μM neuron pairs (black; ^**^P<0.01; Log-rank Mantel-Cox test).

To test whether depolarization block of SST interneurons directly contributes to behavioral seizures, we generated mice in which the GqDREADD excitatory chemogenetic receptor was specifically expressed in SST interneurons (SST-Cre; GqDREADD^+/-^, Figure 4A). GqDREADD receptors were activated using CNO (i.p.) at 0.2, 1, and 5 mg/kg and changes in ECoG activity monitored (Figure 4A). In control mice (SST-Cre; GqDREADD^-/-^), CNO administration at 5 mg/kg did not alter the ECoG or induce seizure behavior (Figure 4B). In contrast, SST-Cre; GqDREADD^+/-^ mice exhibited a robust hypersynchronization of ECoG activity characterized by an increase in low-frequency (2-10Hz) power and displayed behavioral manifestations associated with *status epilepticus* including loss of balance, and clonic jerks (Figure 4C; Video 5). The fact that the effect of CNO administration in SST-Cre; GqDREADD^+/-^ mice was dose-dependent (^*^P<0.05 comparing SST-Cre;GqDREADD^-/-^; black; n=4 and SST-Cre;GqDREADD^+/-^; red; n=8; unpaired t-test; Figure 4D) taken alongside the absence of effect of CNO in SST-Cre;GqDREADD^-/-^ mice supports our notion that GqDREADD-mediated activation of SST interneurons is sufficient to induce prolonged seizures (i.e. *status epilepticus*) in mice expressing wild-type sodium channels. In support of our hypothesis, recordings from SST interneurons in SST-Cre;GqDREADD^+/-^ mice demonstrated an increase in spontaneous excitability (^**^P<0.01; n=8,4 mice; paired t-test; Figure 4E-F) and premature depolarization block in response to depolarizing current injections (^***^P<0.001; n=8,4 mice; paired t-test; Figure 4G-H) after bath application of CNO (10 μM). These findings thus resembled the physiological phenotype observed in mutant *SCN8A*-expressing SST interneurons (Figure 2).

**Figure 4.**
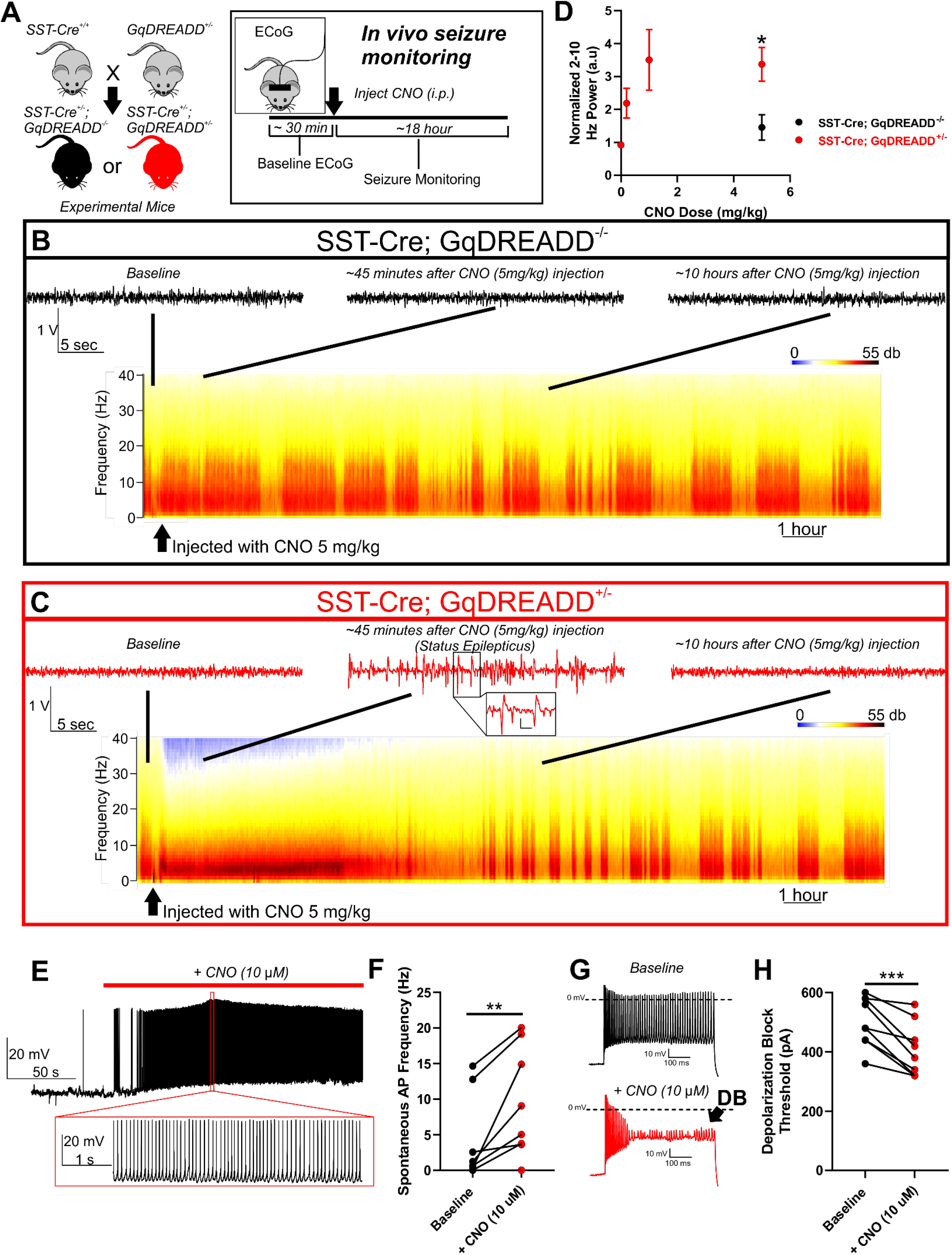
Chemogenetic activation of SST interneurons in WT mice is sufficient to induce seizures. **A**. Breeding strategy: Female mice homozygous for SST-cre (SST-cre^+/+^) were bred with male mice heterozygous for a floxed GqDREADD allele (GqDREADD^+/-^) to produce offspring that were either SST-Cre^+/-^; GqDREADD^-/-^ control mice (black) or SST-Cre^+/-^; GqDREADD^+/-^ experimental mice (red). For *in vivo* seizure monitoring, ECoG was recorded from each mouse for ∼30 minutes of baseline activity, then treated with vehicle or CNO (i.p. at 0.2, 1, and 5 mg/kg), and monitored for seizure activity for ∼18 hours. **B-C**. Example relative power spectra of ECoG activity for ∼18 hours with representative ECoG traces before CNO injection (baseline), ∼45 minutes after CNO injection, and ∼10 hours after CNO injection. **B**. Example traces indicating that in mice lacking the GqDREADD receptor (black), CNO treatment (5 mg/kg) did not lead to seizure behavior or changes in ECoG activity. **C**. Example traces reveal that upon CNO administration GqDREADD^+/-^ mice (red) exhibited highly synchronized ECoG activity and spike-wave discharges indicative of *status epilepticus* which recovers within ∼8 hours. Inset shows expanded view of example spike-wave discharges. Scale bar is 0.25 volts and 0.2 sec. **D**. Normalized 2-10 Hz power relative to pre-injection baseline observed in response to vehicle and CNO (0.2, 1, and 5 mg/kg). SST-Cre; GqDREADD^+/-^ mice (n=5 for vehicle, 0.2, and 1 mg/kg, n=8 for 5 mg/kg; red) experienced a robust and dose-dependent increase in 2-10 Hz power. SST-Cre; GqDREADD^-/-^ control mice (black) did not show a significant increase in 2-10 Hz power upon administration of 5 mg/kg CNO (N=4). **E**. Representative example trace of an SST-Cre; GqDREADD^+/-^ SST interneuron showing spontaneous increase in excitability in response to bath application of CNO (10 μM; red bar). Expanded view illustrates the high firing frequency during CNO exposure. **F**. Spontaneous firing frequencies of SST interneurons from SST-Cre; GqDREADD^+/-^ mice before (black) and after (red) treatment with CNO (10 μM; n=8, 4 mice; ^**^P<0.01; paired t-test). **G**. Example traces of SST interneuron excitability from an SST-Cre; GqDREADD^+/-^ mouse in response to a 600 pA current injection before (black), and after (red) CNO (10 μM) bath application. Premature depolarization block (DB, arrow) is observed after CNO treatment. **H**. Depolarization block threshold for each SST-Cre; GqDREADD^+/-^ SST interneuron before (black) and after (red) treatment with CNO (10 μM; n=8, 4 mice; ^***^P<0.001; paired t-test).

The most prominent biophysical impairment of both the N1768D and R1872W voltage-gated sodium channel variants is an elevated steady-state persistent sodium current (I_NaP_) (Baker et al., 2018; Bunton-Stasyshyn et al., 2019; Lopez-Santiago et al., 2017; Ottolini et al., 2017; Veeramah et al., 2012; Wengert et al., 2019a). We recorded I_NaP_ in SST interneurons using slow voltage-ramps (Figure 5A). I_NaP_ from *Scn8a*^*D*/+^ (63 ± 8 pA; n=13, 4 mice) and *Scn8a*-SST^W/+^ (68 ± 8 pA; n=15, 3 mice) SST interneurons were augmented relative to WT interneurons (38 ± 4 pA; n=12, 4 mice; ^*^P<0.05 ANOVA followed by Dunnett’s multiple comparisons test; Figure 3B-E). Half-maximal voltages (V_1/2_) were not different between groups (Figure 3F). The transient sodium current was recorded in excised somatic patches in order to assess activation and steady-state inactivation properties under proper voltage-control (Figure 5G). No differences in current density, voltage-dependent activation, or steady-state inactivation between WT (n=9,4 mice), *Scn8a*^*D*/+^ (n=13,5 mice) and *Scn8a*-SST^W/+^ (n=8, 3 mice) SST interneurons were detected (Figure 5H-M; Table 2). Together, these results suggest that an increased I_NaP_ current magnitude is likely the primary cause for the altered excitability observed in the mutant SST interneurons and subsequent entry into depolarization block.

**Table 2:** Voltage-gated sodium channel parameters from somatic outside-out recordings WT, *Scn8a*^*D*/+^, and *Scn8a*-SST^W/+^ somatostatin inhibitory interneurons.

|  | Peak Current (pA) | Activation V<sub>1/2</sub> (mV) | Activation Boltzmann Slope Factor | Inactivation V<sub>1/2</sub> (mV) | Inactivation Boltzmann Slope Factor | Decay Tau (ms) |
| --- | --- | --- | --- | --- | --- | --- |
| WT (n=13) | -505 ± 117 | -38.7 ± 2.2 | 5.1 ± 0.4 | -67.9 ± 2.7 | 9.6 ± 0.9 | 0.54 ± 0.04 |
| *Scn8a*<sup>D/+</sup> (n=9) | -451 ± 87 | -37.7 ± 1.6 | 5.9 ± 0.6 | -65.0 ± 3.1 | 10.1 ± 0.7 | 0.66 ± 0.05 |
| *Scn8a*-SST<sup>W/+</sup> (n=8) | -498 ± 153 | -40.4 ± 2.9 | 5.9 ± 0.8 | -65.8 ± 2.8 | 11.2 ± 1.0 | 0.68 ± 0.05 |

Values are mean ± SEM. All comparisons with WT were non-significant using a one-way ANOVA.

**Figure 5.**
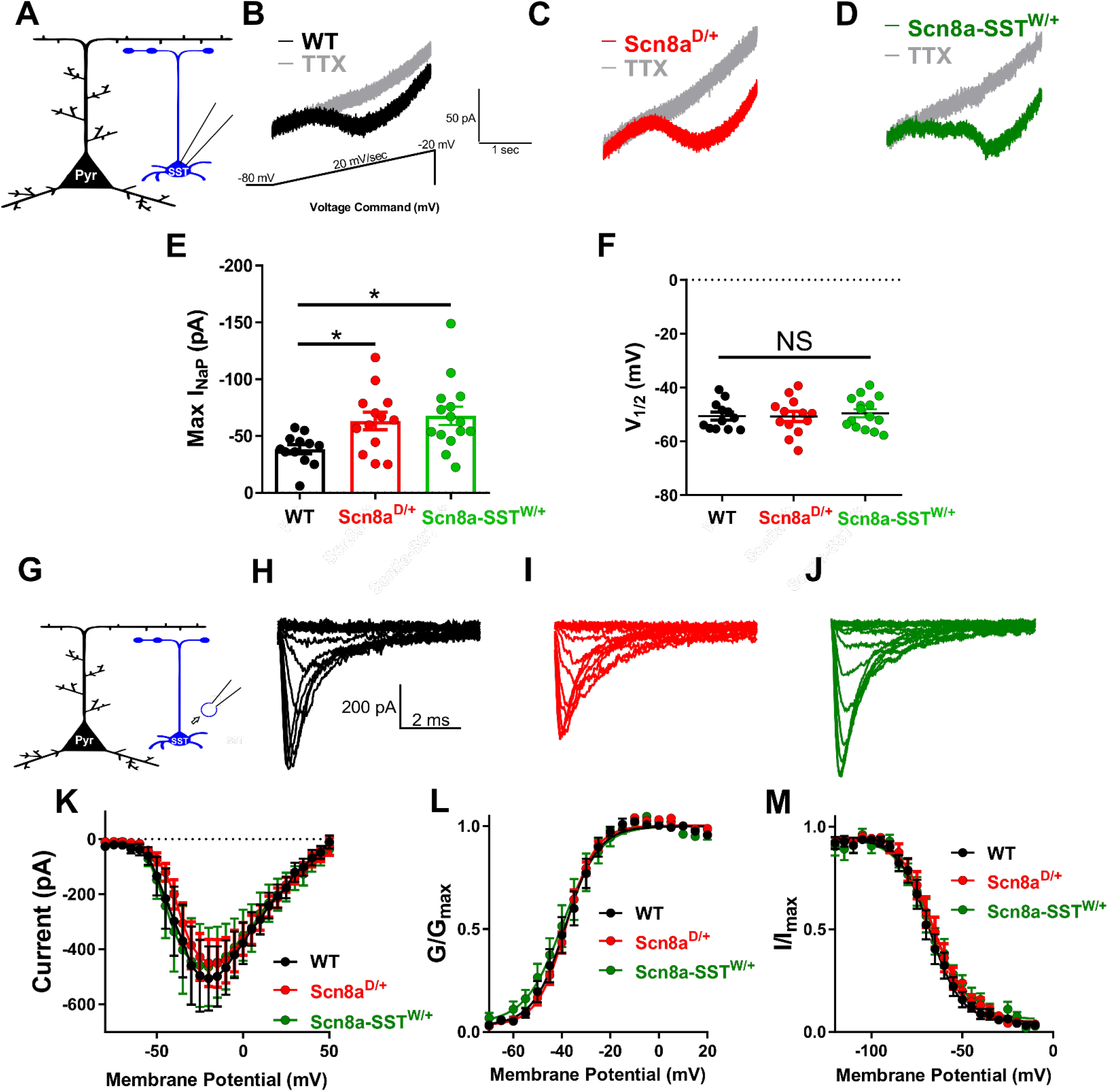
Elevated steady-state persistent sodium currents in *Scn8a*^*D*/+^ and *Scn8a*-SST^W/+^ somatostatin interneurons. **A**. Whole-cell recordings were collected from SST interneurons (blue) to measure whole-cell persistent sodium currents. **B-D**. Example traces of steady-state persistent sodium currents evoked by slow voltage ramps (−80 mV to −20 mV at 20 mV/sec) before (black, red, or green) and after addition of TTX (500 nM; gray) for WT (B; black), *Scn8a*^*D*/+^ (C; red), and *Scn8a*-SST^W/+^ (D; green) SST interneurons. **E**. Elevated maximum persistent sodium current (I_NaP_) in *Scn8a*^*D*/+^ SST interneurons (red; n=13, 4; ^*^P<0.05) and *Scn8a*-SST^W/+^ (green; n=14, 3; ^*^P<0.05) compared to WT SST interneurons (black; n=12, 4; One-way ANOVA followed by Dunnett’s multiple comparisons test). **F**. Half-maximal voltage of activation between WT, *Scn8a*^*D*/+^, and *Scn8a*-SST^W/+^ SST interneurons was not significantly different between groups (NS; P>0.05). **G**. Somatic transient sodium current was assessed in SST interneurons (blue) using patch-clamp recordings in the outside-out configuration. **H-J**. Example traces for family of voltage-dependent sodium currents recorded from outside-out excised patches from either WT (H; black), *Scn8a*^*D*/+^ (I; red), and *Scn8a*-SST^W/+^ (J; green) SST interneurons. **K**. Current-voltage relationship for WT (black; n=9, 4 mice), *Scn8a*^*D*/+^ (red; n=13, 5 mice), and *Scn8a*-SST^W/+^ (green; n=8, 3 mice) SST interneurons. **L-M**. Voltage-dependent conductance and steady-state inactivation curves for WT (black), *Scn8a*^*D*/+^ (red), and *Scn8a*-SST^W/+^ (green) SST interneurons.

To test our hypothesis, we utilized a single-compartment conductance-based computational neuron model to examine neuronal frequency-current relationships as we varied the magnitude of the persistent sodium conductance (g_NaP_). At low levels of g_NaP_, neuronal firing frequency increased with the injected current magnitude over a large range of depolarizing current injections, reaching a maximum of 96 APs/500-ms (Figure 6A-C). In support of our hypothesis, we found that increasing the magnitude of the g_NaP_ led to a decrease in rheobase current and a corresponding facilitation of AP firing at low magnitude current injections, followed by premature AP failure via entry into depolarization block as the current injection magnitude was increased (Figure 6A-C). Although the neuronal model used is overly simple in both geometric and ionic terms relative to SST interneurons *in vivo*, the sufficiency of the model to reproduce both features of mutant SST interneuron excitability is strong evidence that an elevated I_NaP_ is directly sufficient for initial hyperexcitability followed by premature entry into depolarization block.

**Figure 6.**
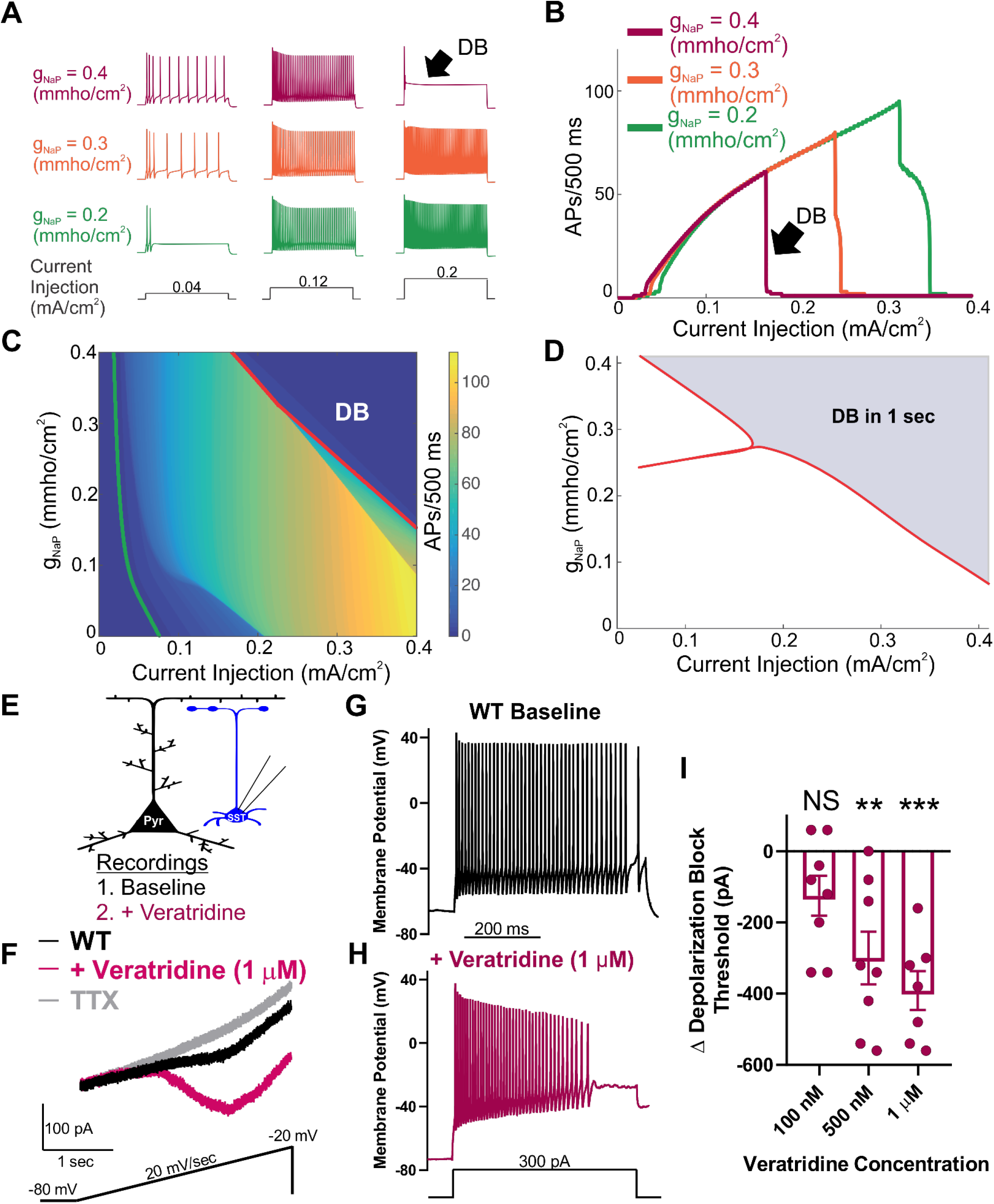
Augmentation of the persistent sodium current induces depolarization block. **A**. Example traces of action potentials elicited in response to 500 ms current injections in an *in silico* neuronal model. The maximal persistent sodium conductance was varied from g_NaP_ = 0.2 mmho/cm^2^ (green), through g_NaP_ = 0.3 mmho/cm^2^ (orange) to g_NaP_ = 0.4 mmho/cm^2^ (magenta), whilst the magnitude of the current injection was increased from 0.04 mA/cm^2^ (left), through 0.12 mA/cm^2^ (middle) to 0.2 mA/cm^2^ (right). Increasing g_NaP_ increased the number of APs observed over the stimulation period at low current magnitudes, and led to failure of AP initiation at higher current injection magnitudes (black arrow, DB). **B**. Number of action potentials observed over the 500-ms stimulus relative to the current magnitude. As g_NaP_ increases, the current magnitude required to induce depolarization block (black arrow, DB) decreases. **C**. Heat map showing the number of APs as both g_NaP_ and the current injection magnitude are varied. Also shown is the rheobase current required to induce at least one AP (green line) and the curve at which the neuron ceases to generate repetitive APs as it enters depolarization block (red line, DB). **D**. Two parameter bifurcation showing the location of dynamic transitions leading to depolarization block as the g_NaP_ and the current magnitude are varied. The red line demarcates the shaded region of parameter space in which depolarization block occurs (over a 1s current pulse) from the unshaded region in which it does not. Further information about the bifurcations can be found in the Supplementary Information. **E**. Experimental design: Whole-cell recordings were made from an SST interneuron (blue) before (baseline) and after bath application of the I_NaP_ current activator, veratridine (magenta). **F**. Example trace of I_NaP_ current before (black) and after application of veratridine (1 μM; magenta). **G-H**. Example traces of a WT SST interneuron before (G; black) and after (H; magenta) bath application of I_NaP_ current-activator veratridine (1 μM). **I**. Shift in depolarization block threshold in response to bath-applied veratridine at 100 nM (n=8, 3 mice), 500 nM (n=8, 3 mice), and 1 μM (n=7, 3 mice). Significance determined by paired t-test (^**^P<0.01; ^***^P<0.001).

Transitions in dynamical states, such as the entry into depolarization block can be understood through the application of bifurcation theory. This approach allowed for tracking of the number and properties of steady states (corresponding to the neuron being at rest or in depolarization block) and periodic orbits (corresponding to spiking behavior) as model parameters are varied. Providing further evidence for our hypothesis, we observed that, as g_NaP_ was increased, the depolarization block boundary shifted leftwards to lower current magnitudes, corroborating our simulation results and providing a more detailed theoretical underpinning for them (Figure 6D). Of note, our results indicate that the dynamical transition into depolarization block results from the interplay between persistent sodium currents and slow potassium currents (detailed description provided in Supplementary Information).

To test predictions from the computational model regarding the effect of increasing persistent sodium conductance, we collected recordings of WT SST interneurons before and after application of veratridine. Veratridine is a plant-derived toxin that increases the magnitude of the I_NaP_ (Alkadhi and Tian, 1996; Bikson et al., 2003; Mantegazza et al., 1998; Otoom and Alkadhi, 2000; Tazerart et al., 2008; Tsuruyama et al., 2013). In agreement with our hypothesis, veratridine (1 μM) increased the SST interneuron I_NaP_ by 68 ± 20 pA (n=6, 2 mice; ^*^P<0.05; Paired t-test; Figure 6F). In current-clamp recordings, we compared the depolarization block threshold before and after treatment with veratridine at 100 nM, 500 nM, and 1 μM. We observed a dose-dependent reduction in depolarization block threshold in response to veratridine at 500 nM (^**^P<0.01; Paired t-test; n=8, 3 mice) and 1 μM (^***^P<0.01; Paired t-test; n=7, 3 mice), but not at 100 nM (n=8, 3 mice; Paired t-test; Figure 6G-I). Taken together, our computational and pharmacological evidence demonstrate that the magnitude of the I_NaP_ current strongly influences the threshold for depolarization block in SST interneurons.

## Discussion

In this study, we have identified a mechanism by which SST interneurons are impaired and contribute to seizures in *SCN8A* encephalopathy. We show that 1) expression of the *SCN8A* mutation R1872W selectively in SST interneurons is sufficient to induce audiogenic seizures, implicating a previously unidentified role for SST interneurons in *SCN8A* encephalopathy, 2) gain-of-function *SCN8A* mutations facilitate SST interneuron intrinsic hyperexcitability and AP failure via depolarization block, 3) rhythmic instances of depolarization block of mutant SST interneurons are coincident with cortical pyramidal neuron ictal discharge under seizure-like conditions, 4) chemogenetic activation and induction of depolarization block of WT SST interneurons induces prolonged seizures, further supporting a critical role for SST interneurons in maintaining balanced neuronal network excitation, and 5) the I_NaP_ is elevated in SST interneurons expressing gain-of– function *SCN8A* mutations, and it directly facilitates depolarization block. These findings provide novel insight into a previously unappreciated role for SST interneurons in seizure initiation, not only in the context of *SCN8A* encephalopathy, but to epilepsy more generally.

Previous studies using transgenic mice models of *SCN8A* encephalopathy have focused primarily on the impact of gain-of-function *SCN8A* mutations on excitatory neurons and how pro-excitatory alterations in sodium channel properties render these neurons hyperexcitable and corresponding networks seizure prone (Baker et al., 2018; Bunton-Stasyshyn et al., 2019; Lopez-Santiago et al., 2017; Ottolini et al., 2017; Wengert et al., 2019). Our finding that cell-type-specific expression of a patient-derived gain-of-function *SCN8A* mutation (R1872W) in SST interneurons, but not forebrain excitatory neurons, resulted in audiogenic seizures indicates that this neuronal population contributes to seizures in *SCN8A* encephalopathy.

A proconvulsant impact of inhibitory interneuron hypofunction has been reported previously in the context of Dravet syndrome in which expression of loss-of-function *SCN1A* results in reduced intrinsic excitability of inhibitory interneurons (Cheah et al., 2012; Escayg and Goldin, 2010; Rhodes et al., 2004; Tai et al., 2014). In contrast, our recordings in SST interneurons expressing gain-of-function *SCN8A* mutations exhibited an initial hyperexcitability followed by progressive AP failure due to depolarization block. Interestingly, our results were similar to those found in a recent study examining the gain-of-function *SCN1A* point-mutation T226M which leads to hypofunction through depolarization block and clinically results in a distinct epileptic encephalopathy more severe than traditional Dravet Syndrome (Berecki et al., 2019; Sadleir et al., 2017). Our findings support their interpretation that gain-of-function voltage-gated sodium channel variants can produce functionally dominant-negative effects through depolarization block-mediated AP failure.

Making up 5-10% of cortical neurons, SST inhibitory interneurons are critical components of cortical microcircuits (Nigro et al., 2018; Rudy et al., 2011). In cortical layer V, SST interneurons have been shown to provide lateral inhibition to local pyramidal neurons upon high-frequency AP firing (Silberberg and Markram, 2007). This disynaptic inhibitory mechanism has been suggested to regulate the overall excitability of the cortical column, preventing simultaneous excitability in many pyramidal neurons at one time (Silberberg and Markram, 2007). In our dual-recording experiments, in which an SST interneuron and a nearby pyramidal neuron were recorded simultaneously under seizure-like conditions, we observed that SST interneuron depolarization block events and pyramidal neuron ictal discharges were coincident. Depolarization block of SST interneurons could alter the excitability of nearby pyramidal neurons by transiently silencing synaptic inhibition from SST interneurons which have been shown to act in concert to strongly constrain the excitability of cortical pyramidal neurons (Safari et al., 2017). An additional, non-mutually exclusive mechanism, is that depolarization block in SST interneurons could change the local extracellular ionic environment in a manner that elevates pyramidal neuron excitability. In a previous modeling study of interplay between interneurons and pyramidal neurons during *in vitro* seizure-like events, elevations in external potassium concentration due to interneuron depolarization block was a critical parameter in achieving good agreement between the model and experimental results (Ullah and Schiff, 2010).

An interplay between inhibitory and excitatory neurons via depolarization block of inhibitory interneurons and network seizure-like activity has been previously proposed (Cammarota et al., 2013; Parrish et al., 2019; Swadlow, 2003; Ziburkus et al., 2006). In the study by Parrish *et al*, depolarization block of PV, but not SST interneurons occurred during seizure-like events *in vitro*, although recording in the cell-attached configuration might prohibit one to definitively detect the brief instances of depolarization block in SST interneurons observed in our study (Parrish et al., 2019). To our knowledge, our study is the first to show that cortical SST-interneurons, similar to other fast-spiking interneurons, have coordinated activity with nearby pyramidal neurons characterized by coincident depolarization block events during pyramidal ictal discharges.

Our chemogenetic experiments further support our hypothesis that hypersensitivity to depolarization block of SST interneurons directly contributes to seizures *in vivo*. GqDREADD-mediated hyperactivation of WT SST interneurons increased spontaneous excitability and increased susceptibility to depolarization block (similar to the effects of mutant *SCN8A* expression) and remarkably, led to prolonged seizure activity and behavior (i.e., *status epilepticus*). These surprising results indicate that aberrant SST physiology, namely increased propensity for depolarization block, is sufficient to induce hypersynchronous ECoG activity and seizures in otherwise normal mice. Further experiments are warranted to continue to clarify the role that SST interneurons have in various types of epilepsy and whether SST depolarization block is pathogenic in other contexts.

We observed a profound increase in the magnitude of the I_NaP_ in both *Scn8a*^*D*/+^ and *Scn8a*-SST^W/+^ SST interneurons. The I_NaP_ has been intensely investigated since its discovery (French et al., 1990; Stafstrom, 2007; Stafstrom et al., 1985; Wengert and Patel, 2020). It has been ascribed a variety of important physiological and neurocomputational functions including spike timing (Osorio et al., 2010; Vervaeke et al., 2006), amplification of synaptic inputs (Schwindt and Crill, 1995; Stuart, 1999; Stuart and Sakmann, 1995), and pacemaking (Del Negro, 2002; Yamada-Hanff and Bean, 2013; Yamanishi et al., 2018; Zhong et al., 2007). An aberrantly large I_NaP_ has also been implicated in various epilepsy syndromes (Hargus et al., 2011; Lossin et al., 2003; Rhodes et al., 2004; Stafstrom, 2007; Veeramah et al., 2012; Wengert and Patel, 2020). In our recordings of mutant SST interneurons, we found higher rates of spontaneous activity, depolarized resting membrane potential, decreased rheobase, hyperpolarized AP threshold, altered upstroke and downstroke velocity, and elevated input resistance, each of which has been previously associated with an augmented I_NaP_ (Ceballos et al., 2017; Herzog et al., 2001; Stafstrom et al., 1982; Yamada-Hanff and Bean, 2013). Moreover, by modifying the magnitude of the I_NaP_ alone, our computational model recapitulated both of the primary physiological features of the mutant interneurons, that of initial hyperexcitability at low current injections followed by premature depolarization block at higher current injections. Our findings that veratridine, a toxin previously utilized to interrogate the mechanism of the I_NaP_ (Alkadhi and Tian, 1996; Bikson et al., 2003; Mantegazza et al., 1998; Otoom and Alkadhi, 2000; Tazerart et al., 2008; Tsuruyama et al., 2013) increased the I_NaP_ and induced premature depolarization block in WT SST interneurons, support our computational modeling. Further, our bifurcation analysis revealed that the dynamic transitions from resting to spiking to depolarization block were each dependent upon the I_NaP_ magnitude, indicating that the I_NaP_ magnitude is of general importance for determining neuronal spiking dynamics. Thus, an increased I_NaP_, regardless of particular mechanism, would be predicted to induce hypersensitivity to depolarization block.

Within the class of SST interneurons there is a great deal of diversity with respect to morphology, gene-expression, synaptic input and output, and physiology. In our recordings of cortical layer V SST interneurons we likely recorded from numerous distinct subpopulations of inhibitory interneurons including Martinotti cells, basket cells, bitufted, horizontal, and multipolar cells (Yavorska and Wehr, 2016). Ongoing efforts to distinguish subcellular populations within somatostatin-positive interneurons will further refine the interpretation of the results presented here.

### Conclusions

Previous studies have contributed significant evidence to the notion that *SCN8A* epileptic encephalopathy is caused by gain-of-function *SCN8A* mutations which result in hyperexcitability of excitatory neurons and renders the network hyperexcitable and seizure-prone. In this report, we have demonstrated that dysfunctional SST inhibitory interneurons also contribute to *SCN8A* epileptic encephalopathy: An elevated I_NaP_ in SST inhibitory interneurons results in hyperexcitability and AP failure due to depolarization block. Gain-of-function *SCN8A* mutations render SST interneurons more sensitive to depolarization block, and under seizure conditions, these instances of SST depolarization block are coincident to pyramidal neuron ictal discharges.

Induction of SST interneuron hyperexcitability and depolarization block, even in the absence of an *SCN8A* mutation, paradoxically leads to *status epilepticus* illustrating the critical contribution of SST interneurons to seizures. Restoration of normal excitability in SST inhibitory interneurons may provide a novel therapeutic strategy for patients with *SCN8A* encephalopathy.

## Methods

### Ethics Approval

All procedures were conducted in accordance with University of Virginia’s ACUC guidelines.

### Mouse Husbandry and genotyping

*Scn8a*^*D*/+^ and *Scn8a*^*W*/+^ were generated as previously described and maintained through crosses with C57BL/6J mice (Jax#: 000664) (Bunton-Stasyshyn et al., 2019; Wagnon et al., 2015). Cell-type specific expression of R1872W was achieved using males homozygous or heterozygous for the R1872W allele and females homozygous for EIIa-Cre (Jax#: 003724), EMX1-Cre (Jax#: 005628), or SST-Cre (Jax#:013044) to generate mutant mice (*Scn8a*-EIIa^W/+^, *Scn8a*-EMX1^W/+^, and *Scn8a*-SST^W/+^ respectively) (Bunton-Stasyshyn et al., 2019). Control mice for audiogenic experiments in *Scn8a*-EIIa^W/+^ and *Scn8a*-EMX1^W/+^ were separate litters lacking the R1872W allele and for *Scn8a*-SST^W/+^ they were littermate controls lacking the R1872W allele. Fluorescent labeling of SST interneurons was achieved by first crossing male *Scn8a*^*D*/+^ or *Scn8a*^*W*/+^ mice with a Cre-dependent tdTomato reporter (Jax#:007909) and then with female mice homozygous for SST-Cre. Experimental groups used roughly equal numbers of male and female mice to control for any potential sex differences. For chemogenetic experiments, male mice heterozygous for GqDREADD allele (Jax#: 026220) were crossed with female mice homozygous for SST-Cre. All genotyping was conducted through using Transnetyx automated genotyping PCR services.

### Audiogenic Seizure Test

To test for audiogenic seizures mice were taken from their home cage and transferred to a clean test cage where they were allowed to acclimate for ∼20 seconds before the onset of the acoustic stimulus. Similar to a method described previously (Martin et al., 2020), a sonicator (Branson 200 ultrasonic cleaner) was used to produce the audiogenic stimulus directly adjacent to the test cage. The stimulus duration lasted for 50 seconds or until the animal had a behavioral seizure. *Scn8a*-EIIa^W/+^ mice were tested between P13-P16. *Scn8a*-Emx1^W/+^ were tested between 21-24 days. *Scn8a*-SST^W/+^ mice were tested at 7-10 weeks. Videos of audiogenic seizure tests were recorded with a laptop webcam and the duration of seizure phases was analyzed by taking the time in seconds that the mouse spent in each of the phases: a wild-running phase characterized by fast circular running throughout the cage, a tonic phase characterized by hindlimb extension and muscle rigidity, a clonic phase typified by myoclonic jerking of the hindlimbs, and recovery exemplified when the mouse ceased myoclonic jerking and righted itself.

### Brain Slice Preparation

Preparation of acute brain slices for patch-clamp electrophysiology experiments was modified from standard protocols previously described (Baker et al., 2018; Bunton-Stasyshyn et al., 2019; Wengert et al., 2019a). Mice were anesthetized with isoflurane and decapitated. The brains were rapidly removed and kept in chilled ACSF (0°C) containing in mM: 125 NaCl, 2.5 KCl, 1.25 NaH_2_PO_4_, 2 CaCl_2_, 1 MgCl_2_, 0.5 L-Ascorbic acid, 10 glucose, 25 NaHCO_3_, and 2 Na-pyruvate. The slices were constantly oxygenated with 95% O_2_ and 5% CO_2_ throughout the preparation. 300 μM coronal or horizontal brain sections were prepared using a Leica VT1200 vibratome. Slices were collected and placed in ACSF warmed to 37°C for ∼30 minutes then kept at room temperature for up to 6 hours.

### Electrophysiology Recordings

Brain slices were placed in a chamber continuously superfused (∼2 mL/min) with continuously oxygenated recording solution warmed to 32 ± 1 ^o^C. Cortical layer V SST interneurons were identified as red fluorescent cells and pyramidal neurons were identified based on absence of fluorescence and pyramidal morphology via video microscopy using a Zeiss Axioscope microscope. Whole-cell recordings were performed using a Multiclamp 700B amplifier with signals digitized by a Digidata 1322A digitizer. Currents were amplified, low-pass filtered at 2 kHz, and sampled at 100 kHz. Borosilicate electrodes were fabricated using a Brown-Flaming puller (Model P1000, Sutter Instruments) to have pipette resistances between 1.5-3.5 MΩ.

### Intrinsic Excitability Recordings

Current-clamp recordings of neuronal excitability were collected in ACSF solution identical to that used for preparation of brain slices. The internal solution contained in mM: 120 K-gluconate, 10 NaCl, 2 MgCl_2_, 0.5 K_2_EGTA, 10 HEPES, 4 Na_2_ATP, 0.3 NaGTP (pH 7.2; osmolarity 290 mOsm). Intrinsic excitability was assessed using methods adapted from those previously described (Ottolini et al., 2017; Wengert et al., 2019a). Briefly, resting membrane potential was manually recorded immediately going whole-cell and confirmed using a 1-min gap-free recording of the neuron at rest. Current ramps from 0 to 400 pA over 4 seconds were used to calculate passive membrane and AP properties including threshold as the point at which membrane potential slope reaches 5% of the maximum slope, upstroke and downstroke velocity which are the maximum and minimum slopes on the AP respectively, the amplitude which was defined as the voltage range between AP peak and threshold, the APD_50_ which is the duration of the AP at the midpoint between threshold and peak, input resistance which was calculated using a −20 pA pulse in current clamp recordings, and the rheobase which was operationally defined as the maximum amount of depolarizing current that could be injected into the neurons before eliciting APs. AP frequency-current relationships were determined using 500-ms current injections ranging from −140 to 600 pA. Action potentials were counted only if the peak of the action potential was greater than 0 mV. Threshold for depolarization block was defined in this context as the current injection that resulted in the maximum number of APs, which for neurons that did not undergo depolarization block was the maximum current injection 600 pA.

### Outside-Out Voltage-gated Sodium Channel Recordings

Patch-clamp recordings in the outside-out configuration were collected using a protocol modified from an approach previously described (Ottolini et al., 2017). The recording solution was identical to that used for brain-slice preparation. The internal solution for all voltage-gated sodium channel recordings contained in mM: 140 CsF, 2 MgCl_2_, 1 EGTA, 10 HEPES, 4 Na_2_ATP, and 0.3 NaGTP with the pH adjusted to 7.3 and osmolality to 300 mOsm. Voltage-dependent activation and steady-state inactivation parameters were recorded using voltage protocols previously described (Wengert et al., 2019a).

### Persistent Sodium Current Recordings

The recording solution was modified with a reduced sodium concentration to allow for proper voltage control through the entire range of command voltages (Royeck et al., 2008; Wengert et al., 2019b). It contained in mM: 50 NaCl, 90 TEACl, 10 HEPES, 3.5 KCl, 2 CaCl_2_, 2 MgCl_2_, 0.2 CdCl_2_, 4 4-AP, 25 D-glucose. Steady-state persistent sodium currents were elicited using a voltage ramp (20 mV/sec) from −80 to −20 mV. After collecting recordings at baseline, the procedure was repeated in the presence of 1uM tetrodotoxin (TTX) to completely isolate the sodium current. TTX-subtracted traces were analyzed by extracting the current at each mV from −80 mV to −20 mV. The half-maximal voltage for activation of the current was calculated as previously described (Wengert et al., 2019a). Any recordings in which the neuron escaped steady-state voltage control were discarded before TTX application.

### Dual-cell Recordings

Gap-free recordings of pairs of one pyramidal and one SST interneuron <200 μM away were used to examine cell interplay during seizure-like events. Seizure-like events were evoked using Mg^2+^-free ACSF containing either 50 or 100 μM 4-aminopyridine (4-AP). Seizure-like depolarization block events were defined in this context as instances in which the membrane potential reached a stable value above action potential threshold which occurred simultaneously with burst of action potentials in the nearby pyramidal neuron.

### Computational Modeling

A single-compartment conductance-based neuronal model was modified based on one used previously (Nowacki et al., 2012). The dynamics of the neuronal voltage is described by:

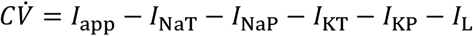

where *V* is the transmembrane voltage, *C* is the membrane capacitance, *I*_app_ is the applied current density, *I*_NaT/NaP_ are the transient and persistent sodium current densities, respectively, and *I*_KT/KP_ are the transient and persistent potassium currents densities, respectively. These currents are specified by the functional forms:

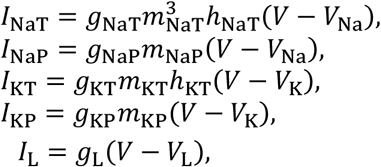

where *g*_X_ represents the maximal conductance of channel type *X* ∈{NaT, NaP, KT, KP}, *m*_X_. represents the fraction of open activation gates of channel type X, with *h*_X_ corresponding to the fraction of open inactivation gates, where appropriate, and *V*_Y_ is the reversal (Nernst) potential of ionic species Y ∈{Na, K, L}. The dynamics of the gating variables obey:

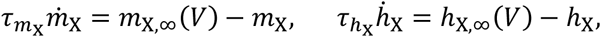

where 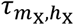 are the timescales of the activation and inactivation of channel type X, respectively; and *m*_*X,∞*_(*V*) and *h*_*X,∞*_(*V*) are its steady state activation and inactivation curves (e.g., Figure 3F-G), which are themselves described by Boltzmann functions of the form:

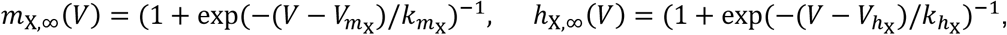

where 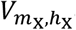 are the thresholds for activation and inactivation and 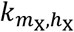 are the associated sensitivities around this threshold. The activation of both types of sodium channel is assumed to be instantaneous, meaning that *m*_NaT,NaP_ *= m*_NaT,NaP,*∞*_(*V*) always. The parameter units and values are as follows:

| Parameter | Unit | Value | Parameter | Unit | Value |
| --- | --- | --- | --- | --- | --- |
| $g_{\text{NaT}}$ | mmho/cm<sup>2</sup> | 130 | $V_{\text{Na}}$ | mV | 60 |
| $g_{\text{NaP}}$ | mmho/cm<sup>2</sup> | [0-0.4] | $V_{\text{K}}$ | mV | -85 |
| $g_{\text{KT}}$ | mmho/cm<sup>2</sup> | 10 | $V_{\text{L}}$ | mV | -65 |
| $g_{\text{KP}}$ | mmho/cm<sup>2</sup> | 1.65 | $\tau_{m_{\text{KT}}}$ | ms | 1 |
| $g_{\text{L}}$ | mmho/cm<sup>2</sup> | 0.02 | $\tau_{m_{\text{KP}}}$ | ms | 75 |
| $V_{m_{\text{NaT}}}$ | mV | -37 | $\tau_{h_{\text{NaT}}}$ | ms | 1 |
| $V_{m_{\text{NaP}}}$ | mV | -47 | $\tau_{h_{\text{KT}}}$ | ms | 1400 |
| $V_{m_{\text{KT}}}$ | mV | -5.8 | $k_{m_{\text{NaT}}}$ | mV | 5 |
| $V_{m_{\text{KP}}}$ | mV | -30 | $k_{m_{\text{NaP}}}$ | mV | 3 |
| $V_{h_{\text{NaT}}}$ | mV | -70 | $k_{m_{\text{KT}}}$ | mV | 11.4 |
| $V_{h_{\text{KT}}}$ | mV | -68 | $k_{m_{\text{KP}}}$ | mV | 10 |
| $I_{\text{app}}$ | mA/cm<sup>2</sup> | [0-0.4] | $k_{h_{\text{NaT}}}$ | mV | -7 |
| $C$ | μF/cm<sup>2</sup> | 1 | $k_{h_{\text{KT}}}$ | mV | -9.7 |

All model simulations were performed in MATLAB R2020a. Code to replicate these simulations is available to download via Git from www.GitHub.com/kyle-wedgwood/DepolarizationBlock.

### Bifurcation Analysis

Bifurcation analysis was performed in DDE-BIFTOOL v3.1.1, which is a MATLAB-based numerical software package for numerical continuation and in AUTO-07P which is a Fortran-based numerical continuation program.

### In-vivo Seizure Monitoring

Custom electrocorticogram (ECoG) headsets (PlasticsOne, Inc.) were implanted in 7–10-week-old SST-Cre^+/-^; GqDREADD^+/-^ or SST-Cre; GqDREADD^-/-^ mice using standard aseptic surgical techniques. Anesthesia was induced with 5% and maintained with 0.5%–3% isoflurane. Adequacy of anesthesia was assessed by lack of toe-pinch reflex. A midline skin incision was made over the skull, and burr holes were made at the lateral/rostral end of both the left and right parietal bones to place EEG leads, and at the interparietal bone for a reference and ground electrodes. A headset was attached to the skull with dental acrylic (Jet Acrylic; Lang Dental). Mice received postoperative analgesia with ketoprofen (5 mg/kg, i.p.) and 0.9 % saline (0.5 ml i.p.) and were allowed to recover a minimum of 2-5 days prior to seizure-monitoring experiments.

Mice were then individually housed in custom-fabricated chambers and monitored for the duration of the experiment. The headsets were attached to a custom low torque swivel cable, allowing mice to move freely in the chamber. ECoG signals were amplified at 2000x and bandpass filtered between 0.3 −100 Hz, with an analogue amplifier (Neurodata Model 12, Grass Instruments Co.). Biosignals were digitized with a Powerlab 16/35 and recorded using LabChart 7 software (AD Instruments, Inc.) at 1 kS/s. Video acquisition was performed by multiplexing four miniature night vision-enabled cameras and then digitizing the video feed with a Dazzle Video Capture Device (Corel, Inc.) and recording at 30 fps with LabChart 7 software in tandem with biosignals.

In seizure monitoring experiments, ∼30 minutes of baseline ECoG signal were recorded before the mice were injected with vehicle or CNO (0.2, 1, and 5 mg/kg). Video ECoG was continuously recorded until the ECoG and behavioral manifestations of status epilepticus ended (typically ∼8 hours). Mice had access to food *ad libitum*, but were restricted from water to protect against headset damage and animal injury during status epilepticus. Power analysis was performed using Spike2 v7.17 (Cambridge Electronic Design, Limited), using FFT size of 1024 points (Hanning method), which provides a 0.9766 Hz resolution across intervals of 1.024 s. To obtain values for the CNO dose-response (Fig. 4D) the peak power in band 2-10 Hz after injection was normalized to pre-injection levels.

### Analysis

All patch-clamp electrophysiology data were analyzed using custom MATLAB scripts and/or ClampFit 10.7. All statistical comparisons were made using the appropriate test using GraphPad Prism 8. One-sided t-tests were only used in the CNO physiology experiments where a clear directional hypothesis was articulated. Data are presented as individual data points and/or mean ± standard error of the mean.

## Supplemental Videos

**Video 1: Audiogenic Seizure in *Scn8a*-EIIa**^**W/+**^ **mouse**.

**Video 2: *Scn8a*-EMX1**^**W/+**^ **mouse does not display audiogenic seizures**.

**Video 3: Audiogenic Seizure in *Scn8a*-SST**^**W/+**^ **mouse**.

**Video 4: WT littermate control does not exhibit sensitivity to audiogenic seizures**.

**Video 5: Behavioral response to CNO (5 mg/kg) in SST-Cre;GqDREADD**^**-/-**^ **control and SST-Cre; GqDREADD**^**+/-**^ **experimental mice**.

## Key Resources Table

| Reagent or Resource | Source | Identifier |
| --- | --- | --- |
| Chemicals |  |  |
| NaCl | Sigma | S7653 |
| KCl | Sigma | P9541 |
| NaH<sub>2</sub>PO<sub>4</sub> | Sigma | S8282 |
| CaCl<sub>2</sub> | Sigma | C3881 |
| MgCl<sub>2</sub> | Sigma | 208337 |
| L-Ascorbic Acid | Sigma | A5960 |
| D-glucose | Sigma | G7021 |
| NaHCO<sub>3</sub> | Sigma | S6297 |
| Na-pyruvate | Sigma | P2256 |
| CsF | Sigma | 198323 |
| EGTA | Sigma | E3889 |
| HEPES | Sigma | H3375 |
| Na<sub>2</sub>ATP | Sigma | A7699 |
| NaGTP | Sigma | G8877 |
| K-gluconate | Sigma | G4500 |
| Veratridine | Sigma | V5754 |
| Clozapine-N-oxide | Sigma | C0832 |
| Tetrodotoxin | Alomone | T-550 |
| Mice |  |  |
| C57BL/6J | The Jackson Laboratory | Stock #: 000664 |
| B6.Cg-Gt(ROSA)26Sortm9(CA G-tdTomatoHze/J | The Jackson Laboratory | Stock #: 007909 |
| SST-Cre | The Jackson Laboratory | Stock #: 013044 |
| EIIA-Cre | The Jackson Laboratory | Stock #: 003724 |
| EMX1-Cre | The Jackson Laboratory | Stock #: 005628 |
| *Scn8a*<sup>D/+</sup> | Gift from Dr. Miriam Meisler |  |
| *Scn8a*<sup>W/+</sup> | Gift from Dr. Miriam Meisler |  |
| B6N; 129-Tg(CAG-CHRM3*, -mCitrine)1Ute/J | The Jackson Laboratory | Stock #: 026220 |
| Software |  |  |
| Prism | GraphPad Software Inc. | https://www.graphpad.com/scientific-software/prism/ |
| MATLAB | MathWorks | https://www.mathworks.com/products/matlab.html |
| pCLAMP Software Suite | Molecular Devices | https://www.moleculardevices.com/products/axon-patch-clamp-system/acquisition-and-analysis-software/pclamp-software-suite |
| LabChart 7 | ADInstrument s, Inc. | https://www.adinstruments.com/support/downloads/windows/labchart-0 |
| Spike2 v7.17 | Cambridge Electronic Design Limited | http://ced.co.uk/downloads/latestsoftware |
| DDE-BIFTOOL v3.1.1 | (Engelborghs et al., 2002) | http://ddebiftool.sourceforge.net/ |
| AUTO-07P |  | https://www.macs.hw.ac.uk/~gabriel/auto07/auto.html |
| Spike |  |  |
| Miscellaneous |  |  |
| Sonicator | Branson Ultrasonics | Model B-200 |

## Supplementary Information

### Bifurcation analysis of hyperexcitability

**Figure S1.**
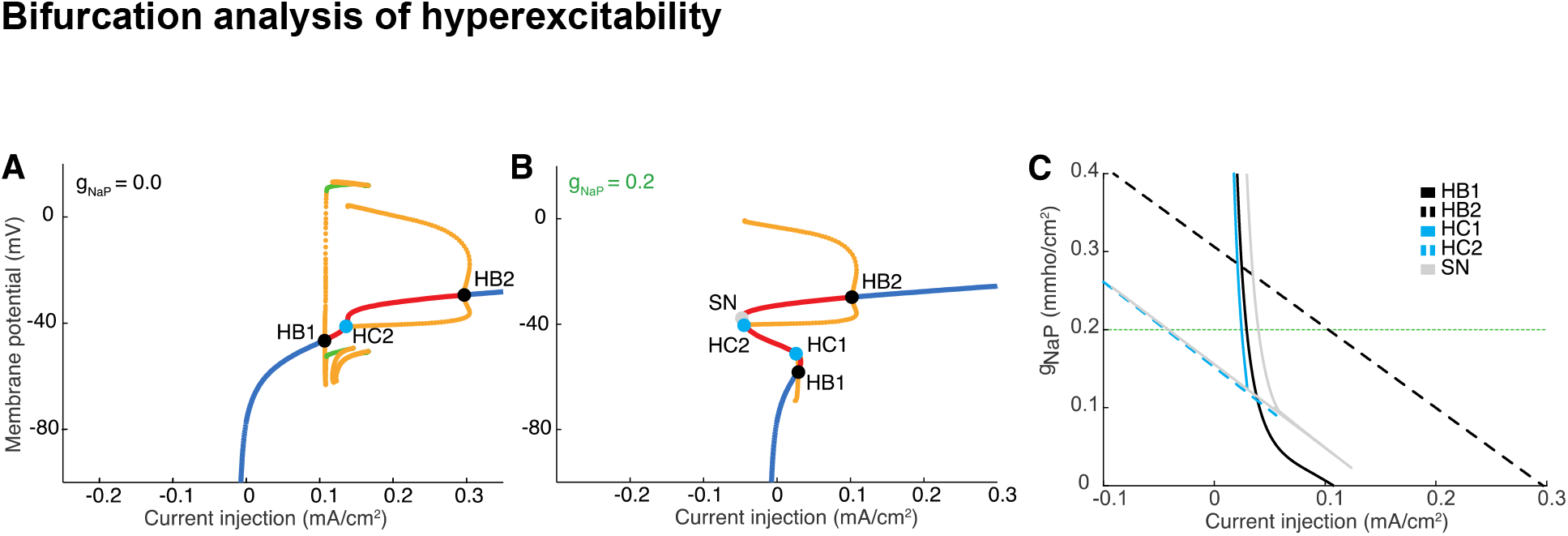
Bifurcation analysis demonstrating hyperexcitability under variation of *g*_NaP_. **A-B:** Bifurcation diagram using *I*_app_ as a control parameter. Here, blue curves represent stable steady states, red curves are unstable steady states and the yellow curves are the maxima and minima of periodic orbits. The transitions from resting to spiking occur via Hopf bifurcations (HB, black dot), whilst saddle node (SN, grey dot) and homoclinic (HC, cyan dot) bifurcations also play a role in shaping overall excitability. In particular, **B** shows that for higher values of *g*_NaP,_ the system supports both resting state behaviour and periodic behaviour over a range of *I*_app_ (between the values of *I*_app_ at which the homoclinic bifurcation, HC2, occurs and that at which the rightmost Hopf bifurcation, HB2, occurs). **C:** Two-parameter continuation in (*I*_*app*_, *g*_*NaP*_) demonstrates that all bifurcation curves exhibit a shift leftwards as *g*_NaP_ increases, highlighting the robustness of the hyperexcitability phenomenon. The horizontal green dotted line indicates the *g*_NaP_ value used in panel B.

To demonstrate hyperexcitability of the neuron as the persistent sodium conductance (*g*_NaP_) is increased, we begin by performing numerical continuation using the magnitude of the applied current (*I*_app_) as a control parameter. This is performed for two distinct values of *g*_NaP_, as indicated in Figure S1A/B. Comparing these figures, we observe that the Hopf and homoclinic bifurcations that give rise to spiking behavior shift to lower values of *I*_app_ as *g*_NaP_ is increased. This means that progressively lower magnitude current injections are required to induce APs, consistent with increased excitability. To confirm this behavior over a range of *g*_NaP_, we then performed, in Figure S1C, two-parameter continuation, allowing *g*_NaP_ and *I*_app_ to vary simultaneously. Here, we see that all bifurcations relevant to transitions into and out of spiking states shift leftwards as *g*_NaP_ is increased.

### The interplay between persistent sodium and transient potassium currents

**Figure S2:**
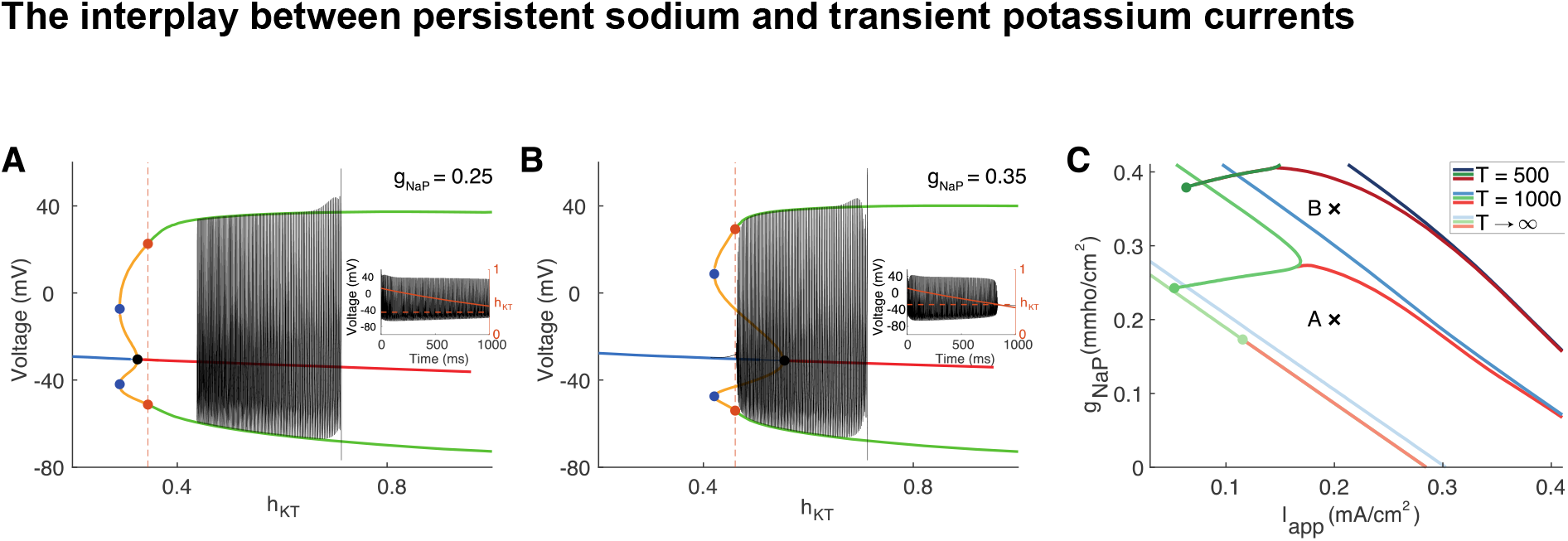
Bifurcation diagrams of the model neuron for fixed *h*_KT._ **A/B:** Bifurcation diagrams of the fast-subsystem obtained by fixing the value of *h*_KT_ and using it as a control parameter for two different values of *g*_NaP_ and setting *I*_app_ = 0.2. Here, the blue (red) curves represent stable (unstable) steady states and the green (orange) represent the maxima and minima of stable (unstable) spiking orbits. Torus bifurcations are marked with orange dots, folds of periodic orbits are marked with blue dots and Hopf bifurcations are marked with black dots. Superimposed on the bifurcation diagram is the trajectory of a simulation in the (*h*_KT,_ *V*) plane. The inset shows the time series of this trajectory, with *V* displayed in black and *h*_KT_ in orange. The value of *h*_KT_ of the rightmost torus bifurcation is indicated by dashed orange lines in both the main panel and the inset. **C:** Two-parameter continuation in (*I*_app,_ *g*_NaP_) of the torus bifurcations (red/orange), period-doubling bifurcations (green) and folds of spiking orbits (blue) of the fast-subsystem subject to the condition that *h*_KT_ at the bifurcation point is equal to that obtained by simulating the slow-subsystem for a duration *T* (with initial conditions given by the steady state values of the system with *I*_app_ = 0). The parameter values used in panels A/B are marked with black crosses.

Examination of model simulations highlights that the transient potassium current plays an important role in determining the entry into depolarization block. In particular, the closure of sufficiently many transient potassium channel inactivation gates (*h*_KT_) marks the entry into the blocked state. Upon noting that the dynamics of *h*_KT_ are significantly slower than all other variables in the system, we may separate the full system into a fast-subsystem and a slow sub-system. For the fast-subsystem, we fix the value of *h*_KT_ and use it as a bifurcation parameter for the other variables. The slow-subsystem describes the evolution of *h*_KT_ in the case where we assume that all other variables are at their *h*_KT_-dependent quasi*-*steady state values i.e. we have

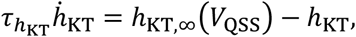

where *V*_*QSS*_ is the solution to

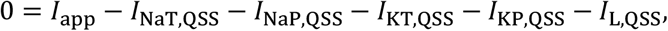

with

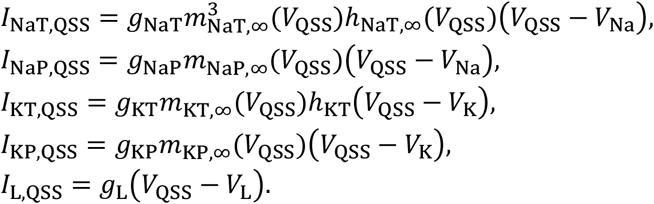

In Figure S2A-B, we plot the bifurcation diagrams for the fast-subsystem for two distinct values of *g*_NaP_. In these figures, we observe the presence of a stable spiking orbit for high values of *h*_KT_. As *h*_KT_ decreases, there is a sequence of instabilities (involving torus and period-doubling bifurcations, depending on the specific parameter values) that destabilise the spiking orbit, which is ultimately terminated in a fold of periodic orbits, after which no periodic solutions exist. A unique steady state exists for all values of *h*_KT_, which is stable for low values of *h*_KT_ and destabilises via a Hopf bifurcation as *h*_KT_ is increased.

We are now in a position to investigate the relationship between *h*_KT_ and depolarization block. At rest, with *I*_app_= 0, *h*_KT_ is high. If we then apply a constant current of sufficiently high magnitude, the neuron will enter the spiking state, after which the value of *h*_KT_ will evolve towards *h*_KT,∞_(*V*_SS_), where *V*_SS_is the steady state of the full system. In the spiking state, the neuron is depolarized compared to its resting state. This fact, in conjunction with the fact that *h*_KT,∞_exhibits a monotonic decreasing dependence on *V*, means that *h*_KT_will decay from its initial value. If the value of *h*_KT_ falls below that associated with any of the bifurcations of the spiking orbit, the neuron will enter depolarization block.

In Figure S2A/B, we see that the first bifurcation of the spiking orbit encountered as *h*_KT_ decreases is a torus bifurcation. The value of *h*_KT_ at this bifurcation is shown by the orange dashed line in the inset, which also shows the trajectories of *h*_KT_ and *V* during the stimulation period. In Figure S2A, the value of *h*_KT_ does not reach the value associated with the torus bifurcation over the stimulation period and so the neuron remains spiking throughout. Contrastingly, in Figure S2B, *h*_KT_ falls below this threshold and the neuron enters the depolarization block state. To demonstrate this behaviour more clearly, the (*V,h*_KT_) components of a model simulation are superimposed on the bifurcation diagrams.

To examine the depolarization block phenomenon over a range of *g*_NaP,_ we perform a two-parameter continuation of the torus, period-doubling and fold of period orbit bifurcations displayed in Figure S2A/B. To address the question of whether the neuron will enter the depolarization block state over a specific duration of stimulus, we need to couple together the fast and slow-subsystems. This is achieved through functionality of DDE-BIFTOOL, which allows for the specification of additional constraints and relationships between parameters. Let us denote by *T* the stimulus duration. We then continue in (*g*_NaP,_ *I*_app_) the bifurcations with the condition that *h*_KT,bif_ = *h*_KT_(*T)*, where *h*_KT,bif_ is the value of *h*_KT_ at the bifurcation and *h*_KT_(*T)* is the solution of the slow-subsystem starting from an initial condition *h*_KT_(0) = *h*_KT,∞_(*V*_rest_), where *V*_rest_ is the resting membrane potential (with *I*_app_ = 0). The solution of the slow-subsystem is found using Matlab’s built-in two-point boundary value problem solver. The results of this procedure for *T =* 500 and *T =* 1000 are shown in Figure S2C, in which we also show asymptotic results for *T* → ∞.

## Acknowledgements

The authors thank the entire Patel lab and others at the University of Virginia for thoughtful feedback on this study and manuscript. KW acknowledges Jennifer Creaser for assistance in the use of AUTO-07P.

